# Skeletal Muscle Remodeling in Immobilized Patients: Determined Using a Parameter Estimation Histomorphometric Approach

**DOI:** 10.1101/2020.06.17.157438

**Authors:** Brent Formosa, Asiri Liyanaarachchi, Samantha Silvers, Domenico L. Gatti, Lars Larsson, Suzan Arslanturk, Bhanu P. Jena

**Affiliations:** Department of Physiology, Wayne State University, Detroit, MI 48201, USA; NanoBioScience Institute, Wayne State University, Detroit, MI 48201, USA; Center for Molecular Medicine & Genetics, Wayne State University, Detroit, MI 48201, USA; Biochemistry, Microbiology and Immunology, School of Medicine, Wayne State University, Detroit, MI 48201, USA; Department of Computer Science, College of Engineering, Wayne State University, Detroit, MI 48201, USA; Department of Physiology and Pharmacology, Karolinska Institutet, SE-171 77 Stockholm, Sweden

**Keywords:** Immobilization-induced myopathy, immunohistochemistry, machine learning, diagnostic pathology

## Abstract

Skeletal muscle biopsy commonly used for light microscopic, electron microscopic and biochemical and transcriptional evaluation remains the gold standard for establishing the etiology of a myopathy. While most myopathies exhibit one or more phenotypes, early stages or several metabolic myopathies often exhibit normal muscle morphology, making diagnosis difficult. In such cases where standard staining techniques fail to offer definitive diagnostic information, a combination of expensive and time-consuming electron microscopy and biochemical testing is required to provide definitive diagnosis. As a step toward overcoming these limitations in diagnostic pathology of skeletal muscle tissue, here we report the application of parameter estimation machine learning approaches on immunofluorescent images of human skeletal muscle tissue acquired using fluorescent microscopy. The machine learning morphometric approach enables the recognition of fine cellular changes in skeletal muscle tissue, allowing determination of skeletal muscle remodeling as a consequence of immobilization.

## Introduction

Alongside non-invasive clinical examination, bioelectrical impedance analysis, electromyography, magnetic resonance spectroscopy, ultrasonography, computed tomography and X-ray absorptiometry (1–7), are among the approaches used to diagnose myopathy. However, skeletal muscle biopsy commonly used for light microscopic, electron microscopic and biochemical and transcriptional evaluation of the skeletal muscle remains the gold standard for establishing the etiology of a myopathy (8–12). Myopathy observed in light and electron micrographs of biopsy tissue include focal myofiber damage as in mitochondrial disorders, segmental damage in dystrophies, or multifocal damage in various inflammatory myopathies (13,14). Furthermore, common myopathic features include variable fiber size with both atrophied and hypertrophied muscle fibers. Atrophied fibers are often rounded, as opposed to the angulated atrophic fibers observed in neurogenic myopathy. Certain myopathies exhibit ragged red fibers which are common in mitochondrial disease. Other myopathies exhibit central core structures, found in central core disease, and the presence of rimmed vacuoles is found in inclusion body myositis (15–17). While most myopathies exhibit one or more such phenotypes, early stage or metabolic myopathies often exhibit normal muscle morphology on routine histochemical examination, making diagnosis difficult. In such cases where standard staining techniques fail to offer definitive diagnostic information, a combination of expensive and time-consuming immunohistochemistry, electron microscopy and biochemical testing is required to provide definitive diagnosis of the disease and for its appropriate treatment and management. To overcome these limitations in diagnostic pathology of skeletal muscle tissue, in the current study, we have developed a parameter estimation histomorphometry approach, that will in the immediate future, be combined with an inexpensive nanoscale imaging approach referred to as differential expansion microscopy (DiExM) (18), to diagnose myopathy. In the current study we use skeletal muscle biopsies obtained on day-1 and again on day-12 from patients following immobilization, to determine changes in muscle morphology, i.e., remodeling of myosin bundles. Utilizing parameter estimation machine learning approach on immunofluorescent images of human skeletal muscle tissue, skeletal muscle remodeling as a consequence of immobilization was determined. This machine learning morphometric approach enables the recognition of fine cellular changes in skeletal muscle tissue, impossible otherwise.

## Materials and Methods

### Human skeletal muscle tissue sections

Human tibialis anterior muscle biopsies obtained from both male and female patients ages 52-84 years on day-1 and again on day 12 following admission to the ICU, was used in the study. In humans, the tibialis anterior muscle has a greater (nearly 80%) proportion of the slow muscle fibers. Tissues were handled, stored and sectioned, according to the published procedures and protocols approved by the institutional review board of Karolinska Institutet, Stockholm, Sweden. Part of each skeletal muscle biopsy was fixed in 4% para formaldehyde (PFA), washed in phosphate buffered saline (PBS), prior to setting the tissue in optimal cutting temperature compound (OCT) block, frozen, and cryosectioned at a thickness of 8-10 μm and placed on poly-L-lysine coated glass slides. Tissue sections on glass slides were handled and stored according to the published procedures and protocols approved by the institutional review board of Wayne State University, prior to immunofluorescent staining and expansion microscopy.

### Immunohistochemistry

Eight to 10 μm 4% PFA fixed human skeletal muscle biopsy tissue sections adhered to poly-L-lysine coated glass slides, were used to perform immunohistochemistry. Tissues sections were hydrated for 48h at 4 °C in PBS pH 7.4. Sections were exposed to 10 mM sodium citrate pH 6.0 for 30 min at 60 °C. Tissue sections were washed six times (2 min per wash) using PBS pH 7.4. at RT. Sections were blocked with MaxBlock blocking medium (Active Motif) containing 0.1% Tween-20 for 2 hours at RT, followed by 6 washes (2 min per wash) in PBS pH 7.4 containing 0.1% Tween-20 at RT. Tissue sections were incubated at 4 °C overnight with the primary-conjugated fluorescent antibody [Slow myosin heavy chain MYH7-Alexa Flour 647 (red); Santa Cruz Biotechnology, Inc], at 1:50 dilution using MaxBind (Active Motif) or 0.5 μg/ml of each antibody. To label the nucleus, sections were incubated with 0.2 μM of 4’,6-diamidino-2-phenylindole (DAPI) (Molecular Probes, Life Technologies, Carlsbad, CA) in PBS pH 7.4, washed 2 times (5 min per wash) with PBS containing 0.1% Tween-20 at RT.

### Light microscopy

Imaging was carried out using a Zeiss apotome imager Z1 fluorescence microscope at 10X and 40X magnifications. For each treatment group, 4 spatially distinct and randomly chosen 2-channel (DAPI 461nm and DsRed 568nm) images were acquired at identical illumination. Each image was extracted with Zeiss apotome imager with extended depth of focus rendered through Zen Imaging Software (Carl Zeiss AG).

### Machine Learning Approach

A machine-learning framework was used to enable the recognition of fine cellular changes in skeletal muscle tissue [Figure 1]. This was accomplished by using the immunomicrographs obtained from human skeletal muscle biopsies obtained from patients on Day 1 and Day 12 of immobilization. Micrographs of nuclei stained using DAPI and muscle fibers immunolabeled using the slow myosin heavy chain conjugated Alexa Flour 647 (red), are fed into a Convolutional Neural Network (CNN) model for object segmentation. During training, the biopsy images were manually annotated as foreground (nuclei stain DAPI and slow twitch myosin) and background (cytoplasm) are used to build a predictive model capable of classifying each pixel on an independent validation set. The model was then tested on the remaining biopsies, to automatically identify the boundaries of each organelle (i.e. object segmentation). The parameters associated with the automatically annotated organelles was then fed into a separate artificial neural network (ANN). The ANN is able to distinguish between D1 and D12 tissue, which are visually indistinguishable.

**Figure 1.**
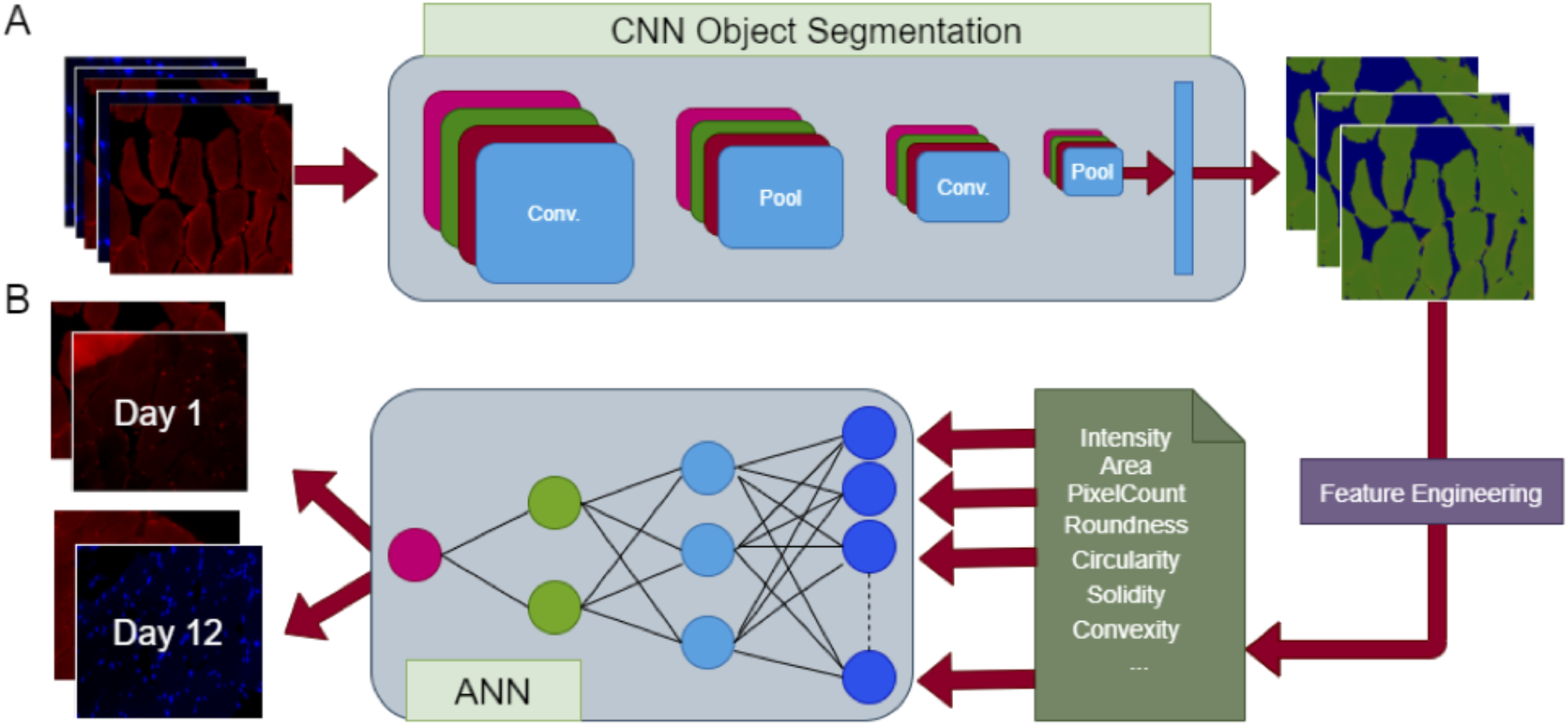
Machine-learning framework enabling the recognition of fine cellular changes in skeletal muscle tissue. **A.** Human skeletal muscle biopsy sections obtained from patients on Day 1 and Day 12 of immobilization, stained for uclei stained with DAPI and fibers immunolabeled using slow myosin heavy chain. Image data are fed into a CNN for object segmentation. During training, images are manually annotated as foreground (nuclei stain DAPI and slow twitch myosin) and background (cytoplasm) and used to build a predictive model capable of classifying each pixel on an independent validation set. The model is then tested on the remaining biopsies to automatically identify the boundaries of each organelle. **B.** Parameters associated with the automatically annotated organelles (e.g. average gray intensity, area, pixel count, etc.) are fed into an ANN to differentiate Day 1 from Day 12 of immobilization.

## Results and Discussion

Immunolabeled human skeletal muscle biopsy obtained from four patients (M639, M623, M626 and M629) on Day 1 (D1, control) and Day 12 (D12, experimental) of immobilization, demonstrate the loss of slow myosin heavy chain immunoreactivity following 12 days of immobilization. However, changes in size and shape of the nuclei stained with DAPI and the myosin bundles between D1 and D12 samples, were visually indistinguishable in all four patients [Figure 2]. To be able to determine the pathophysiological status of the skeletal muscle in ICU patients from immunofluorescent images of human skeletal muscle tissue acquired using fluorescent microscopy, we applied a parameter estimation machine learning approach. The machine learning morphometric approach enables the recognition of fine cellular changes in skeletal muscle tissue visually indistinguishable, allowing determination of skeletal muscle remodeling as a consequence of immobilization.

**Figure 2.**
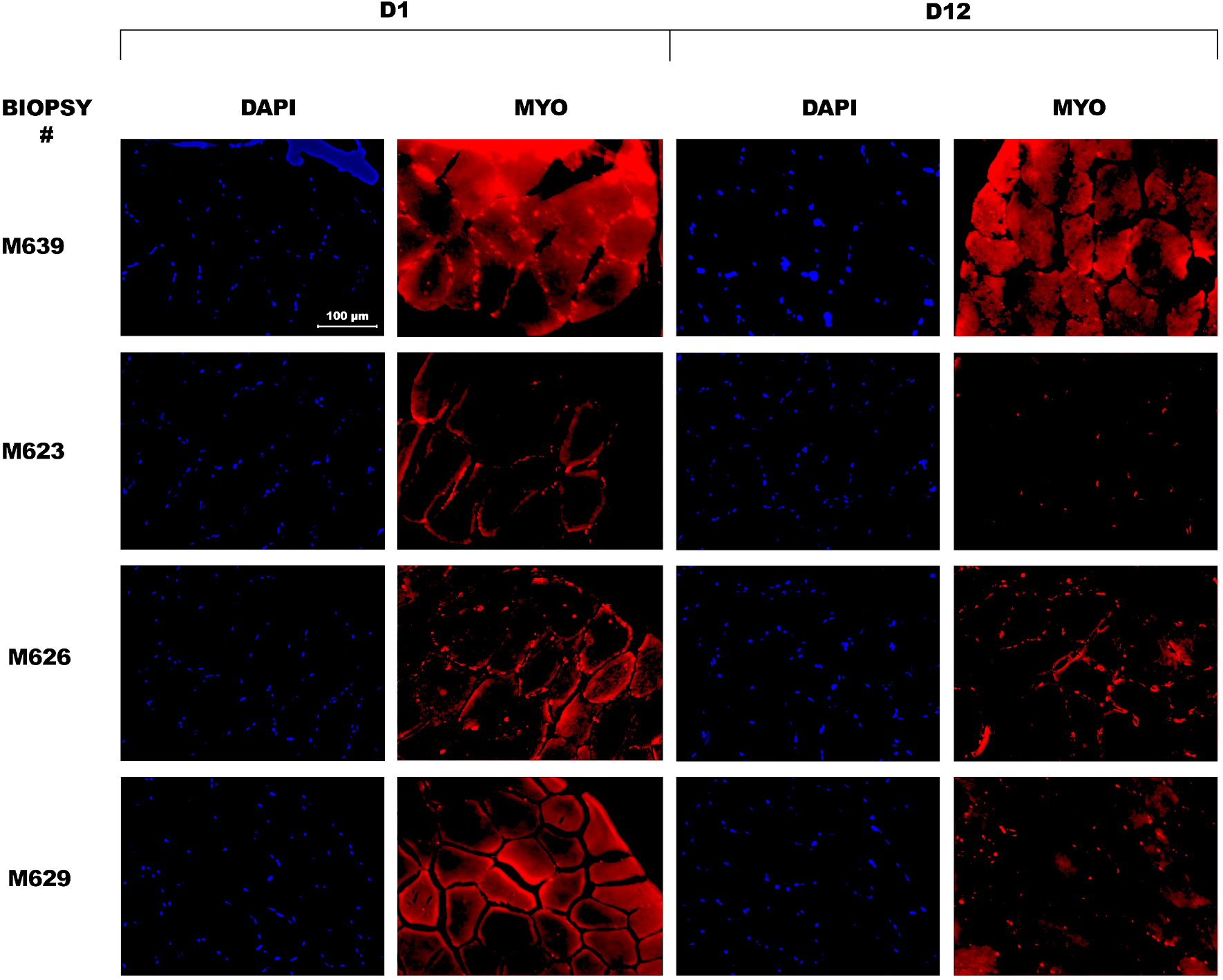
Immunolabeled human skeletal muscle biopsy obtained from four patients (M639, M623, M626 and M629) on Day 1 (D1, control) and Day 12 (D12, experimental) of immobilization using the nuclei stain DAPI and slow myosin heavy chain antibody-conjugated fluorophore (red). Note that there is variability in the amounts and distribution of myosin in different patients in the beginning of the study (D1). Also note the difference between patients in the extent of loss of slow myosin heavy chain immunoreactivity following 12 days of immobilization. However, the size and shape of the nuclei and the myosin bundles between D1 and D12 are visually indistinguishable. *Bar = 100 μm*.

Muscle biopsies fixed with 4% PFA and cryo-sectioned (8-10 um sections), were stained with DAPI, and slow (red) myosin. A convolutional neural network (CNN) was trained using the tissue micrograph data for object segmentation. During training, the sub cellular structures (e.g. nuclei in fibers expressing slow myosin) in multiple images are annotated manually. Once a fully trained CNN model is constructed based on the labeled data, the new biopsy images were fed into the network to automatically annotate each pixel and determine the group of pixels defining each object. To illustrate, Figure 3A shows the human skeletal muscle cells with nuclei stained (blue) and Figure 3B shows the corresponding nuclei segmentation. Similarly, Figure 3D shows the human skeletal muscle biopsy section stained for myosin heavy chain (red) and Figure 3E shows the myosin segmentation using CNNs. The parameters of each organelle are calculated through computational image analysis. Parameters associated with each organelle on day 1 and day 12 are then used as input features fed into a machine learning model to understand significant features associated with prolonged ICU stays. The results of this study serve as a proof of concept in understanding cellular changes in patients with various stages of skeletal muscle myopathy.

**Figure 3.**
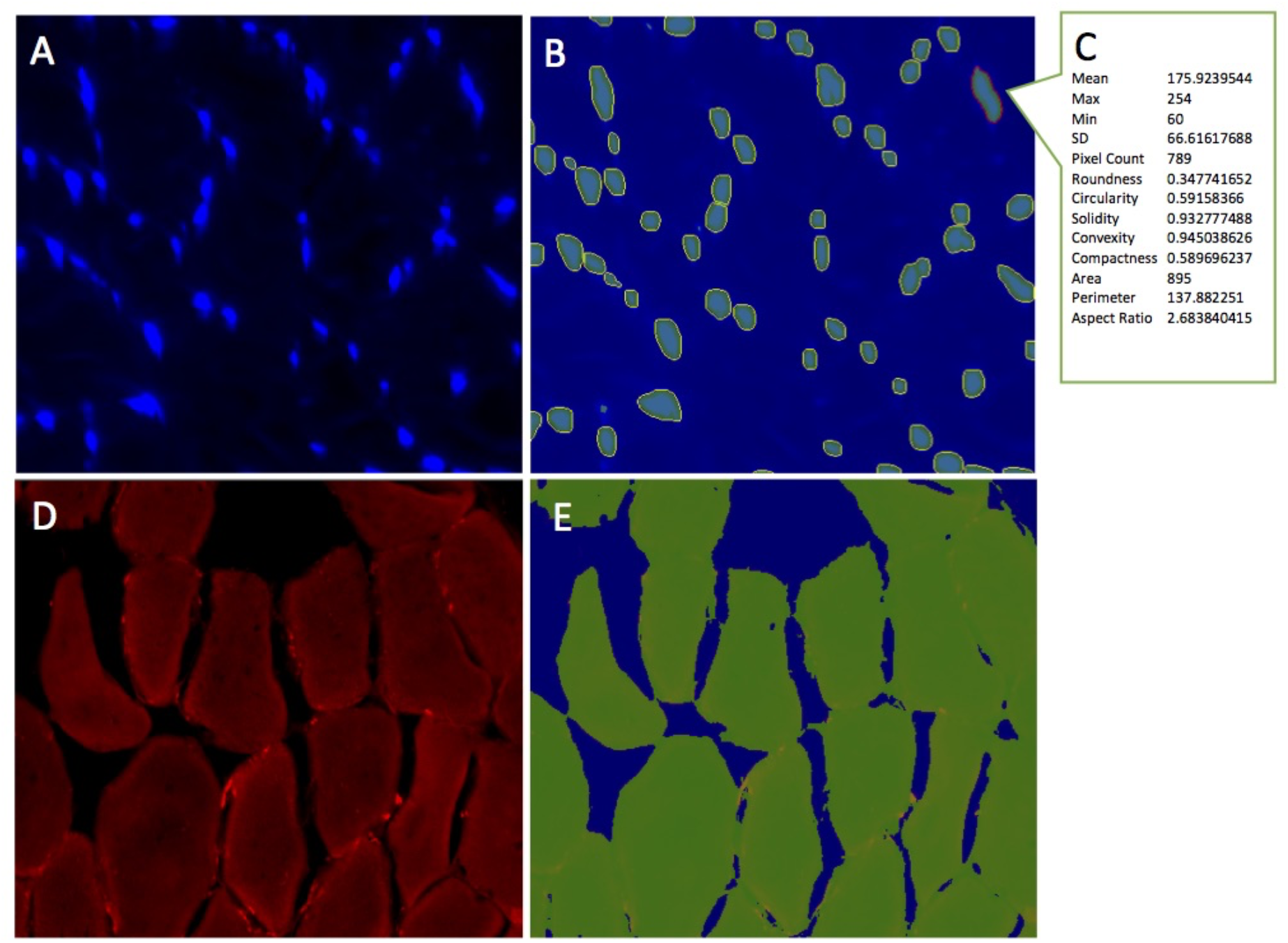
Parameter estimation using a classification model and computational image analysis of human skeletal muscle biopsy tissue. (A) Human skeletal muscle cells with nuclei stained blue (DAPI). (B) Nuclei segmentation and (C) parameter estimation using a classification model and computational image analysis. (D) Human skeletal muscle biopsy section stained for slow myosin heavy chain (E).

An artificial neural network (ANN) is then trained using fluorescent micrographs of D1 and D12 skeletal tissue sections [Figure 2,3] for the identification of significant predictive features, for example those associated with the DAPI-stained nucleus (e.g. area, roundness, circularity, pixel count, convexity etc.), that could differentiate between D1 and D12. at 81% prediction accuracy using a separate validation set. Table I shows the detailed prediction accuracy using DAPI-stained nucleus by the immobilization class.

**Table 1:** Detailed prediction accuracy using DAPI-stained nucleus by immobilization class.

|  | TP Rate | FP Rate | Precision | Recall | F-measure | ROC Area | Class |
| --- | --- | --- | --- | --- | --- | --- | --- |
|  | 0.831 | 0.22 | 0.845 | 0.831 | 0.838 | 0.918 | Day 1 |
|  | 0.78 | 0.169 | 0.762 | 0.78 | 0.771 | 0.918 | Day 12 |
| **Weighted Avg.** | 0.81 | 0.199 | 0.811 | 0.81 | 0.81 | 0.918 |  |

**TP:** True Positive; **FP:** False Positive; **Precision:** Positive Predictive Value; **Recall:** Fraction of the total amount of relevant instances retrieved; **F-measure:** Measure of Tests Accuracy; **ROC Area:** Relative tradeoffs between TP and FP.

Similarly, Table 2 shows the detailed prediction accuracy of the immobilization class using myosin stining. The results have shown that the ANN is able differentiate between D1 and D12 with 88% accuracy. The minimum mean and maximum amount of red intensity are calculated as the most significant predictive features.

**Table 2:** Detailed prediction accuracy of the immobilization class using myosin staining.

|  | TP Rate | FP Rate | Precision | Recall | F-Measure | ROC Area | Class |
| --- | --- | --- | --- | --- | --- | --- | --- |
|  | 0.973 | 0.213 | 0.818 | 0.973 | 0.889 | 0.926 | Day 1 |
|  | 0.787 | 0.027 | 0.967 | 0.787 | 0.868 | 0.926 | Day 12 |
| **Weighted Avg.** | 0.879 | 0.120 | 0.893 | 0.879 | 0.878 | 0.926 |  |

**TP:** True Positive; **FP:** False Positive; **Precision:** Positive Predictive Value; **Recall:** Fraction of the total amount of relevant instances retrieved; **F-measure:** Measure of Tests Accuracy; **ROC Area:** Relative tradeoffs between TP and FP.

In summary, we report in this study the application of parameter estimation machine learning applied on immunofluorescent images of human skeletal muscle tissue, to determine skeletal muscle remodeling as a consequence of immobilization. Besides cellular changes visually indistinguishable in the skeletal muscle micrographs in D1 and D12 of immobilization, there appears to be variability in the amounts and distribution of myosin in different patients in D1 at the beginning of the study. However, this study shows that the machine learning approach utilized, enables the recognition of fine cellular changes visually indistinguishable in skeletal muscle tissue as a consequence of immobilization, allowing determination of skeletal muscle remodeling as a consequence of immobilization. Results of this study serves as a proof of concept in understanding cellular changes in patients with various stages of skeletal muscle wasting, further contributing to the field of diagnostic pathology. Furthermore, the future application of this approach combined with DiExM (18), will help the diagnosis of myopathy at the nano scale.

## ASSOCIATED CONTENT

**Corresponding Author**

## Author Contributions

B.P.J. developed the idea and designed experiments for the study. B.F. and A.L. performed immunofluorescent labeling and prepared samples for differential expansion microscopy. S.A. developed and designed computational and machine learning approaches to assess skeletal muscle remodeling in immobilized patients. B.P.J, L.L, D.L.G. and S.A. wrote the manuscript. L.L. performed studies to obtain cryo-sections of skeletal muscle biopsy tissue from patients on day-1 and from the same patient on day-12 following immobilization. All authors participated in discussions and proofreading the manuscript.

